# Promises and limitations of current ancient human epigenetic clocks

**DOI:** 10.64898/2026.02.04.703756

**Authors:** Youssef Tawfik, Yoan Diekmann, Ludovic Orlando, Joachim Burger, Jens Blöcher

## Abstract

Age-at-death estimation of archaeological human remains is central to palaeodemographic research yet remains particularly challenging for adults where osteological methods often produce imprecise age ranges. Epigenetic clocks can accurately predict chronological age in modern humans, but their applicability to ancient human DNA is unclear due to data limitation and indirect methylation inference. Here, we evaluate the performance of existing epigenetic clocks on reconstructed ancient human methylomes combining high-coverage genomic data and a correction framework adapted to mitigate damage-derived sequence bias. Across multiple CpG window sizes, neither direct clock application nor regression-based retraining produced reliable continuous age-at-death estimates. Reframing age inference as adult-subadult classification did not return statistically supported age classes either. In contrast, sex estimation based on X-chromosome methylation achieved perfect accuracy, before and after correction. Together, these results indicate that current palaeo-epigenetic approaches reliably recover global biological signals but are not sufficiently sensitive to capture gradual, age-related variation in humans. Estimating age-at-death from ancient methylomes will therefore require methodological advances beyond correction alone, including reference data and improved models for inferring damage-derived epigenetic signals.

## INTRODUCTION

Palaeodemography, the study of population dynamics in prehistoric societies, critically depends on accurate age-at-death estimates to reconstruct life-history traits, such as fertility, mortality, and survivorship ^1,2^. However, traditional osteological methods have inherent limitations, both in modern ^3^ and ancient societies ^4^. For subadults, systematic assessment of developmental changes such as tooth eruption and epiphyseal fusion allows for relatively precise age estimates. Once adulthood is reached, skeletal changes become harder to discriminate and correlate only loosely with chronological age, due to high variability in the senescence processes ^5,6^. Furthermore, the “age mimicry” phenomenon, where estimated profiles mirror reference datasets rather than true ages, introduces another layer of systematic bias ^5,7,8^. These limitations restrict the resolution of palaeodemographic models, ultimately limiting our understanding of how ancient populations responded to changes in lifestyle, environmental shifts, cultural transitions, or demographic pressures.

A potential alternative to osteological age estimation has emerged with the development of epigenetic clocks, which represent one of the most significant methodological advances in aging research. Epigenetic clocks started with the discovery that specific patterns of DNA methylation – a central epigenetic mechanism that regulates gene expression, development, and cellular identity ^9,10^ – change predictably with age ^11^. Since then, many epigenetic clocks have been developed across various tissues, based on different modelling approaches ^12–16^. Recent efforts by Horvath and colleagues have also produced universal pan-tissue clocks by focusing on the age-associated methylation changes that are conserved across mammals ^17–20^. Epigenetic clocks rely on the direct quantification of DNA methylation and therefore require high-quality genomic DNA with sequencing depth ∼100X per CpG ^21^, often realized through the use of hybridization-based arrays followed by bisulfite or nanopore sequencing ^22^. This robust methodology, well-established in biomedical and forensic contexts ^23^ could offer a promising framework when extended to ancient DNA (aDNA).

Ancient DNA preserves information about DNA methylation states, either through direct chemical retention of methylated CpG dinucleotides, or indirectly through post-mortem damage patterns that are dependent on original methylation status. Methylated cytosines preferentially deaminate into thymines, while unmethylated cytosines deaminate to uracils ^24^, leaving a molecular signature of past methylation patterns that can be leveraged if specific enzymatic treatments are applied. One such treatment relies on the USER-enzymatic mix, which specifically cleaves uracils, removing post-mortem damage signatures on those deaminated methylated CpGs. These sites are sequenced as CpG dinucleotides when not affected by post-mortem damage, typically outside the short overhanging ends of DNA templates, or as TpGs otherwise, especially at deaminated methylated CpGs ^24^. This discovery established the foundation of palaeo-epigenetics and enabled the reconstruction of the first methylomes of ancient hominins ^25,26^, providing access to regulatory variation not evident from sequencing data of skeletal remains alone ^27^. Several computational frameworks such as epiPALEOMIX ^28^, DamMet ^29^, and RoAM ^30^, utilize these principles to infer methylation levels in ancient genomes.

While applying epigenetic clocks to ancient DNA holds great potential to help advance palaeodemography, and even promises to revive it, direct application has proven to be difficult so far ^25,28,31,32^. However, a recent study on horses introduced a mathematical transformation step adjusting ancient DNA methylation estimates according to modern methylation scales, resulting in improved age-at-death predictions, assuming that CpGs could be covered at high depth-of-coverage ^33^. Following these results, our aim was to investigate whether similar correction strategies can be extended to ancient human genomes and whether a similar increase in precision can be achieved. We therefore compiled a dataset of high coverage genomes (>10X) from individuals with reliable anthropological age-at-death estimates to evaluate the applicability of various epigenetic clocks on ancient humans.

## RESULTS AND DISCUSSION

### Reconstructed ancient methylomes

Our dataset consists of 36 publicly available ancient human genomes (Supplementary Table S1) with corresponding age-at-death estimates (mean anthropological age estimates: 1–65 years) and reported biological sex (11 females and 25 males), each sequenced to high average depth-of-coverage (∼10–34X, see Methods). DNA methylation was reconstructed using DamMet ^29^, which provides probabilistic estimates of the fraction (F) of methylated cytosines at each CpG site. For an accurate assessment of site-specific methylation levels at individual CpGs, substantial sequencing depth is required (>80X, ^29^). To increase precision at lower coverages, CpGs are usually binned within windows of 10–50 CpGs over which estimates are averaged. This approach relies on the strong local correlation of DNA methylation levels among neighboring CpGs that was reported for different tissues ^34^. While this enables estimation of differential methylation for genomes above >10X, it also leads to a degree of smoothing and a reduction of precision at the single site level ^25–29,31^.

As maintaining CpG-level estimation was essential for our downstream analyses, we quantified the decay of correlation among methylation levels of neighboring CpGs with increasing distance using modern whole genome bisulfite sequencing data (WGBS). The strongest correlation could be found within windows of 10–25 CpGs (Supplementary Figure S1). Beyond this range, correlations declined steadily, suggesting that larger windows begin to incorporate CpGs from distinct regulatory elements. Based on our observations we selected windows of 10, 15, 20, and 25 CpGs for downstream analyses. For individual methylomes, the expected bimodal distribution characteristic of mammalian DNA methylation, with enrichment near fully unmethylated (0) and fully methylated (1) states ^35,36^ was observed consistently across all 22 autosomes, thus indicating that the reconstructed F-values reflect true methylation structure rather than random noise (Supplementary Figures S2–S5). CpG islands overlapping with the 40k CpG panel of the mammalian methylation array ^17^ also displayed pronounced hypomethylation (mean ≈ 0.11–0.14), consistent with canonical regulatory signatures observed in modern data for which the sites were selected (Supplementary Figure S6).

We further evaluated whether reconstructed methylomes follow expected inter-individual patterns. A principal component analysis (PCA) on a set of 4,000 X-chromosome linked CpGs ^37^ could clearly separate genetically male and female individuals in our ancient dataset, further demonstrating that the reconstructed methylomes represent biological meaningful features, as shown in Figure 1 for the 10-CpG windows (see Supplementary Figure S7 for results based on windows comprising 15, 20, and 25 CpGs).

**Figure 1.**
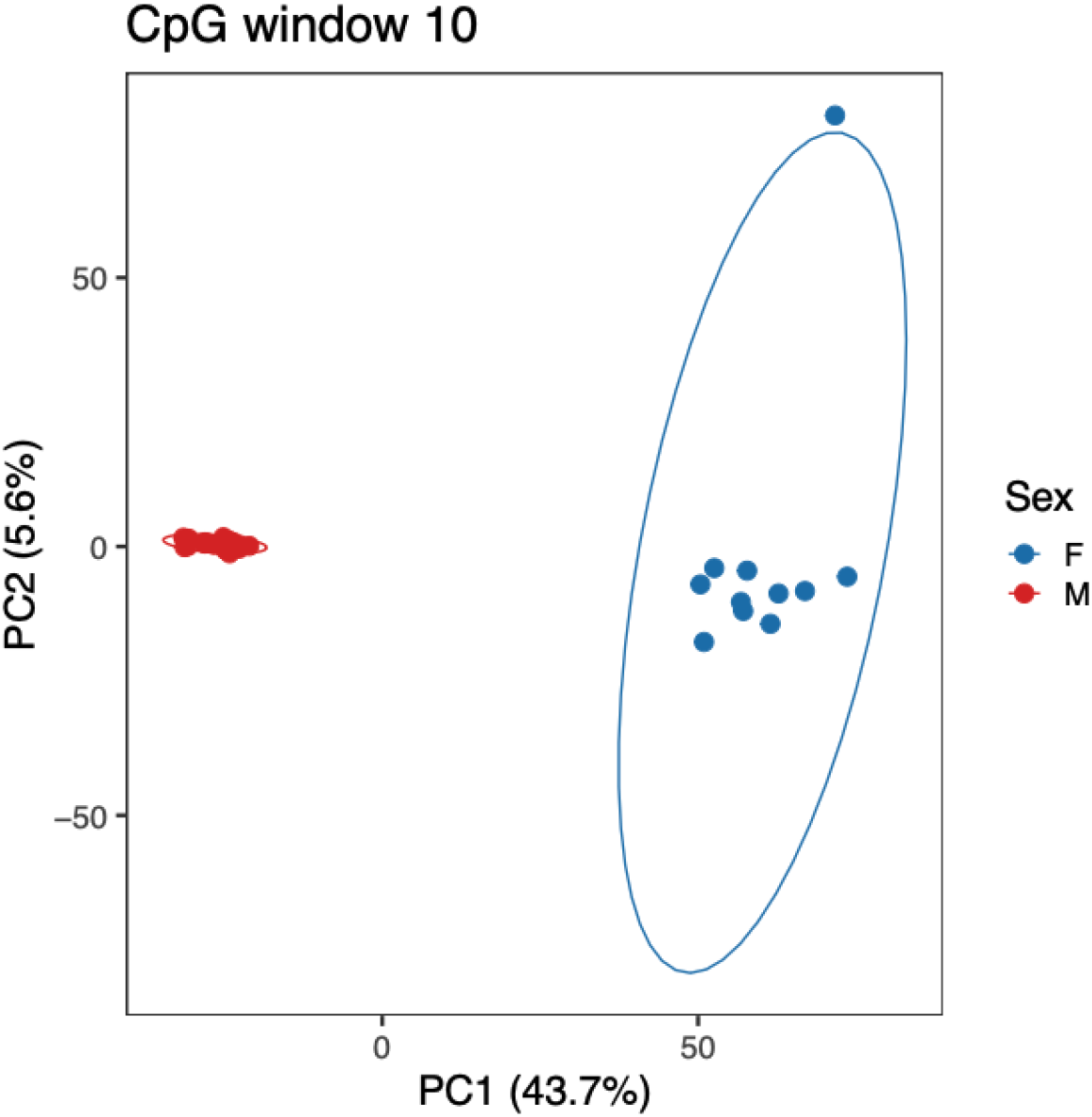
Principal Component Analysis (PCA) of ∼4,000 X-chromosome CpG sites. PC1 (explaining > 40% of variance) clearly separates females (blue) and males (red), capturing sex-specific methylation patterns.

Despite these validations, both previous work ^25,28,31,32^ and our own analyses (Supplementary Figures S8–S11 and Supplementary Table S2) showed that raw DamMet F-values are not directly suitable for age prediction. Applying modern epigenetic clocks to uncorrected F-values resulted in weak associations, with performance varying strongly across CpG window sizes and clock models (Supplementary Figures S11–S14). Notably, these distortions persisted when anthropological age uncertainty was accounted for using interval-aware Monte Carlo analyses, indicating that the unsuccessful observations are not solely driven by imprecision in age-at-death estimates (Supplementary Table S2). Moreover, absolute age estimates were poor and rarely fell within reported anthropological age ranges, indicating a systematic bias between damage-derived F-values and modern DNA methylation levels, necessitating a correction step prior to downstream analyses, as proposed by Liu and colleagues ^33^.

### Correction of reconstructed methylomes

Following Liu *et al*. ^33^, we applied a correction approach, previously introduced for ancient horses, to reduce systematic bias in ancient methylation estimates (F-values) and improve their correlation with modern methylation patterns. Following the correction, Spearman correlation (R) with modern blood-derived array methylomes increased substantially for all CpG window sizes (R=0.57–0.74, 10 CpGs; R=0.60–0.77, 15 CpGs; R=0.61–0.79, 20 CpGs; R=0.61–0.80, 25 CpG; p-value < 0.001 for all, see Figure 2). After correction, PCA on X-linked CpGs still showed the same patterns as prior to correction (Supplementary Figure S12).

**Figure 2.**
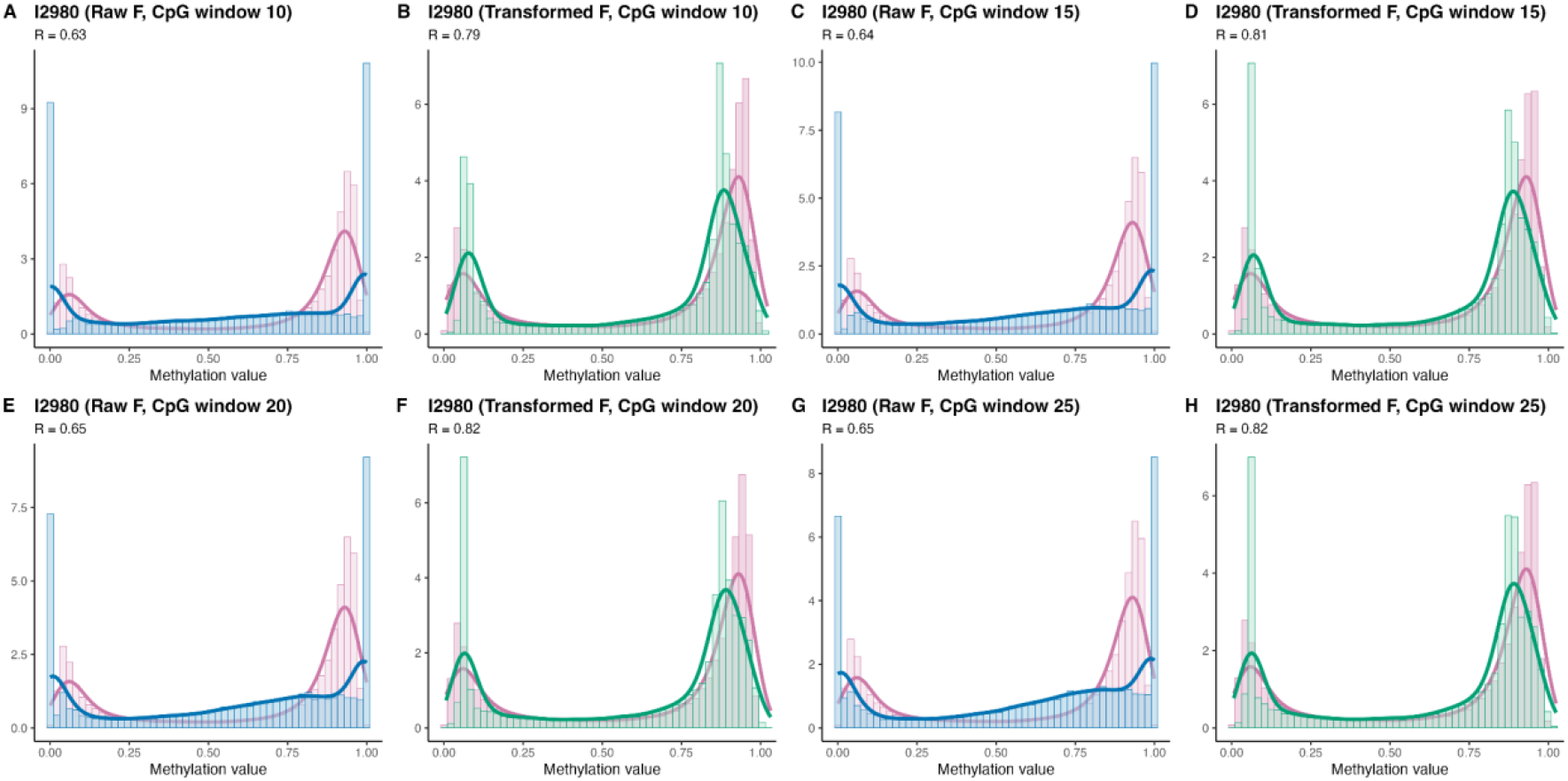
Distribution of methylation values across CpG window sizes before and after correction. Histograms show the distribution of methylation values for sample “I2980”. Overlaid density curves show raw DamMet F-values (blue), transformed methylation (F) values (green), and mean modern methylation values from blood-derived mammalian methylation array data (purple). Across 10, 15, 20, and 25-CpG windows, correction shifts ancient methylation distributions toward that of the modern reference.

Although Spearman correlation was slightly higher in larger CpG windows (for example, from R=0.79 at 10 CpGs to R=0.82 at 25 CpGs in sample I2980 (p-value < 0.001 for both), as shown in Figure 2), these gains were accompanied by only minor changes in overall distributional similarity. Kolmogorov-Smirnov distances differed only marginally across window sizes (0.22, 0.20, 0.19, 0.19, respectively, with p < 0.001 for all), indicating that the improvement from additional smoothing was limited. Therefore, we focused on 10-CpG windows to evaluate whether these corrected methylation value estimates could improve performance in downstream age-at-death prediction models.

### Application of modern epigenetic clocks after correction

To test the direct applicability of established modern epigenetic clocks (with reported mean absolute error of 2-6 years) on the transformed ancient methylomes, four different models were evaluated: (1) the BBT-APM clock ^23^, a tissue-matched model with a small (7) CpG panel; (2&3) two broadly calibrated pan-mammalian/pan-tissue models with large (328 and 793, respectively) CpG panels (UniversalPanMammalianClock1 and UniversalPanMammalianClock2) ^18^, and; (4) the widely used Horvath clock based on age-dependent DNA methylation profiles at 353 CpG sites ^12^.

As anthropological age-at-death is typically interval rather than precise point estimates, conventional point-based error metrics such as mean absolute error (MAE) or root mean square error (RMSE) are not well suited as primary performance measures for our analysis. Instead, individual clock performance was evaluated using an interval-aware Monte Carlo Spearman framework that explicitly accounts for age uncertainty by repeatedly sampling plausible ages within the reported age-at-death ranges and by quantifying the consistency of relative age ordering across ancient individuals (Supplementary Table S3).

Our initial findings for the 10-CpG window (Figure 3) showed that the two universal pan-mammalian clocks exhibited weak but consistent monotonic correlation with anthropological age-at-death, with a median Monte Carlo Spearman (R_MCS_) correlation of 0.33 (95% CI: 0.27–0.39) for UniversalPanMammalianClock1, while UniversalPanMammalianClock2 showed a slightly higher value with 0.36 (95% CI: 0.29–0.42). In contrast, for the BBT-APM clock a consistent negative association with the reported age of –0.18 (95% CI: –0.24 to –0.12) was found, while predictions from the Horvath clock displayed only marginal correlation (0.12; 95% CI: 0.05–0.17).

**Figure 3.**
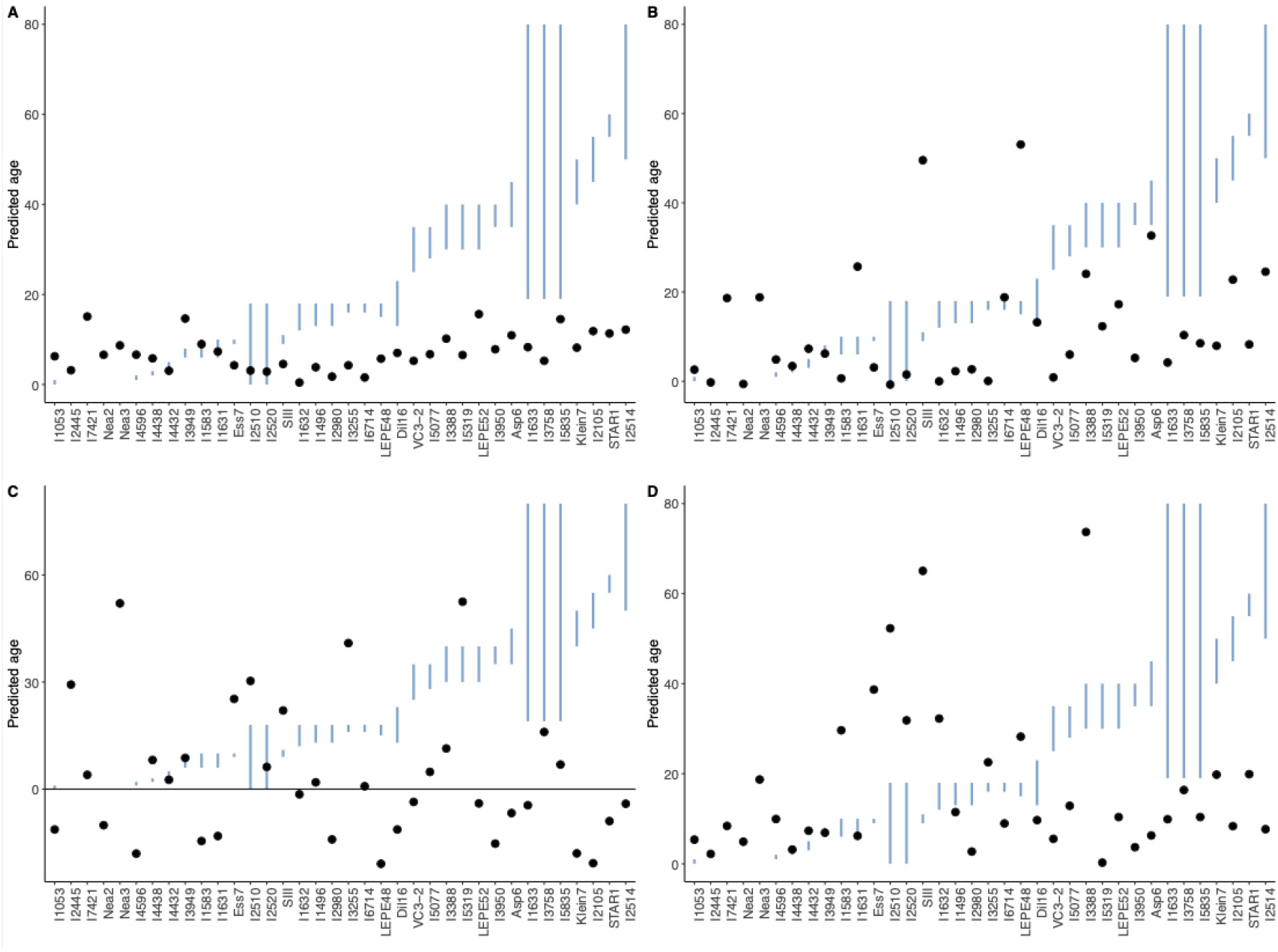
Performance of epigenetic clocks on corrected ancient methylomes (CpG window 10). Panels show results for (A) UniversalPanMammalianClock1, (B) UniversalPanMammalianClock2, (C) BBT-APM, and (D) the Horvath clock. Vertical lines (blue) indicate anthropological age-at-death ranges, and points denote clock-derived predicted ages.

Despite partial preservation of the relative order of ages for some clocks, absolute age estimates were largely inconsistent with anthropological estimates. For 10-CpG windows, only a small fraction of predictions fell within reported age ranges (2.8–13.9%). Moreover, absolute median distances to the nearest anthropological age boundary ranged from approximately 15 years for the two universal pan-mammalian clocks, 20 years for the BBT-APM, and 24 years for the Horvath clock. Thus, even when partial age ordering was recovered, predicted ages were typically implausible in absolute terms, and they deviated substantially from anthropological age ranges. Analyses using larger CpG windows (15, 20, and 25) produced qualitatively similar results (Supplementary Figures S13–S15 and Supplementary Table S3).

Compared to uncorrected analyses, the correction step led to a modest stabilization of age-related rank associations for the pan-mammalian clocks, with higher and more consistently positive Monte Carlo Spearman correlations and narrower confidence intervals. In contrast, correlations derived from raw F-values were generally weaker, less consistent across CpG windows, and in some cases inverted in direction, particularly for the BBT-APM clock (Supplementary Figure S16). Thus, while correction does not recover a strong age signal, it reduces systematic biases present in uncorrected methylation estimates and permits limited, and model dependent, age ordering to emerge.

Given the partial preservation of age-related rank ordering after correction, we next tested whether age-at-death could be recovered by applying a regression model directly on modern blood-derived methylation array data and applying it to reconstructed ancient methylomes. This approach failed to produce meaningful age estimates, with predicted ages being uniformly negative and showing no meaningful association with anthropological age-at-death estimates (Supplementary Figure S17). These results indicate that continuous age regression is not directly transferable to reconstructed ancient methylomes, particularly in the absence of precisely known chronological ages.

Taken together, these results suggest that the primary limitation lies not in the complete absence of age-associated signals, but in the incompatibility between reconstructed ancient methylomes and the assumptions under which the modern epigenetic clocks were trained. Modern clocks are typically developed using precise DNA methylation measurements from fresh tissues ^22^, conditions that differ fundamentally from damage-derived methylation inference in ancient DNA. In addition, the limited precision achievable at a sequencing depths around 10X further impedes reliable age prediction, as accurate methylation inference at the CpG level has been shown to require substantially higher coverage than usually available ^29^. Consequently, reliable age prediction from methylomes might require models explicitly trained on high-precision ancient methylation data, rather than the application of existing clocks or minor adaptation of modern epigenetic clocks. We therefore tested if it is possible to reframe age inference as a binary classification problem rather than a continuous prediction task.

### Adult-subadult classification

While attempts of a direct age-prediction failed so far, we tested if enough age-related information could be retrieved from our reconstructed methylomes to distinguish between broad age classes. We therefore divided our data based on the anthropological age estimates into adults and subadults, using an arbitrary cut-off of 18 years. To further reduce ambiguity in anthropological age-at-death estimates, we restricted the analysis to individuals with clearly defined age ranges, excluding samples with mean estimates overlapping 15–25 years. This resulted in 29 individuals almost evenly distributed among subadults (15) and adults (14). Classification was performed using an elastic net-regularized logistic regression model trained on modern blood-derived methylation array data. When evaluated by cross-validation on the modern dataset, the model achieved perfect classification between the two classes, confirming that this age contrast is readily learnable under conditions of precise age labels and high-quality methylation quantification.

However, when evaluated on our ancient samples, across all CpG window sizes, overall classification accuracy remained limited, with total accuracy never exceeding 60% (Figure 4 and Supplementary Table S4). Apparent high accuracies observed for specific age classes in some models reflected strong classification bias rather than robust age classification, as in the modern dataset. In particular, CpG windows 10 and 25 showed pronounced asymmetry in class assignment, with 10-CpG windows preferentially classifying individuals as adults (adult accuracy 0.79, subadult accuracy 0.33) and 25-CpG windows exhibiting the opposite trend, overclassifying subadults (accuracy for subadults=0.8, and adults=0.36). Although these windows showed modest overall performance (accuracy=0.55 and 0.59, respectively), Fisher’s exact tests indicated no significant association between predicted and anthropological age classes (odds ratios: 1.8 and 2.2, both p > 0.4), showing that apparent class-specific performance was driven by systematic bias rather than reliable age classification. Moreover, 15-CpG windows showed the most balanced performance across age classes, with similar accuracies for adults (0.64) and subadults (0.53), and the highest overall accuracy observed (0.59), yet again not different from a random assignment. The 20-CpG window performed poorly for both classes, with an overall accuracy of only 0.38 and no significant class association as well.

**Figure 4.**
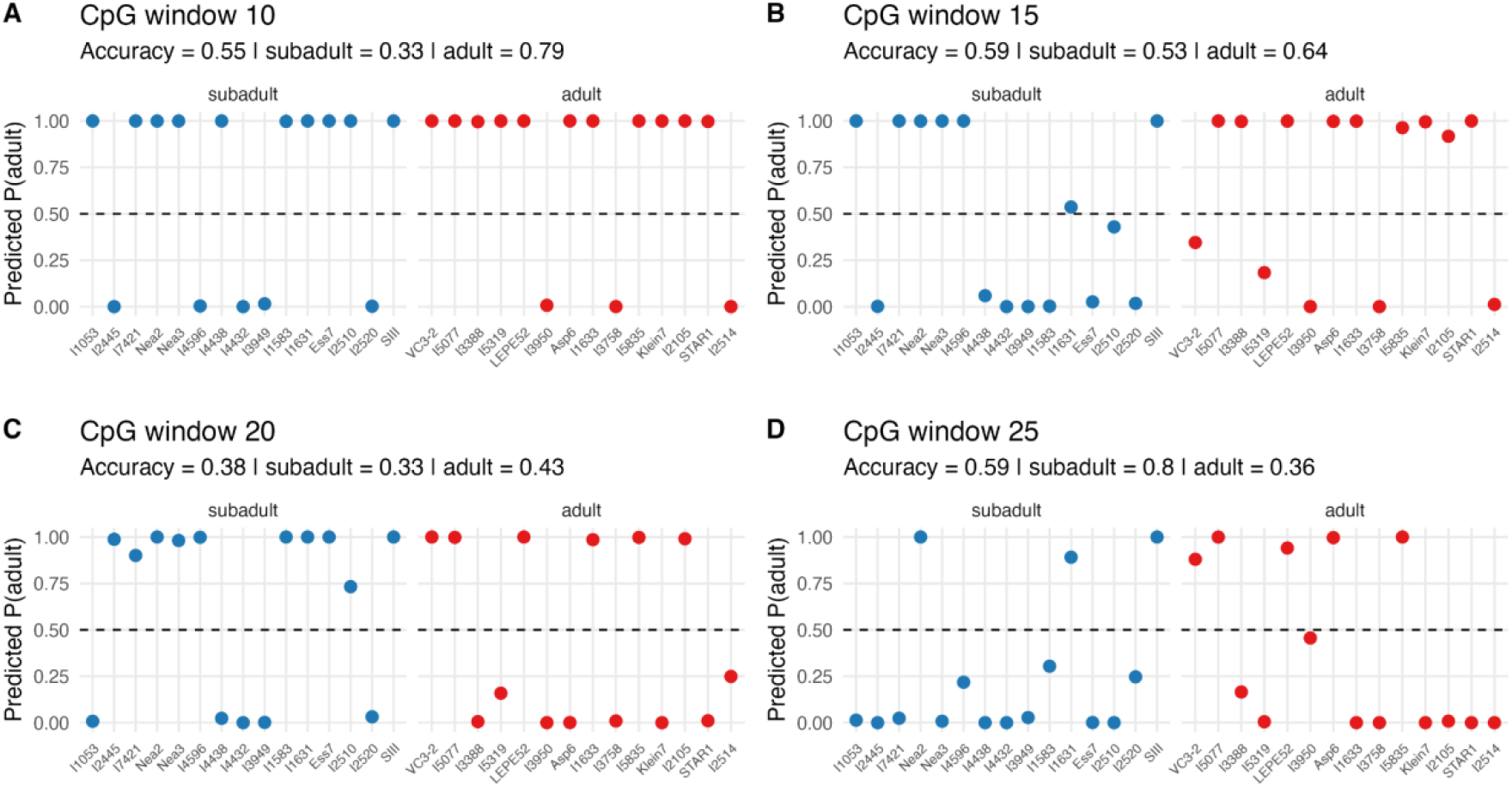
Predicted probability of subadult/adult classification for ancient individuals across different CpG windows. Points represent individual samples, colored by anthropological age classes (subadult; blue, adult; red). Dashed horizontal lines indicate the classification threshold (P = 0.5).

These opposing class-specific error patterns indicate that apparent classification success is largely driven by systematic, window-dependent biases rather than by a stable and biologically meaningful age signal. Consistent with this interpretation, Fisher’s exact tests revealed no significant association between predicted and anthropological age classes for any of the CpG windows, indicating that observed accuracies do not exceed chance expectations. Although results varied across window sizes, none yielded statistically supported age classification.

We further tested whether incorporating ancient samples into model training could stabilize performance, by training mixed models combining modern and ancient individuals. Model evaluation was assessed through *k*-fold cross-validation, in which folds were defined exclusively on the ancient samples. In line with previous results, this approach did not lead to consistent performance improvement and instead showed instability across folds and window sizes, reinforcing the conclusion that limited ancient sample size combined with heterogeneous methylation signal properties largely constrains reliable supervised learning (Supplementary Table S5).

### Conclusions and limitations of this study

This study highlights both the promise and current limitations of ancient epigenetic age estimation when applied to ancient human genomes. By systematically evaluating reconstructed ancient methylomes before and after applying a correction framework adapted from Liu et al. (2023), we show that correction improves similarity between ancient and modern methylation data and allows limited age-related structure to be recovered. However, in contrast to results reported for modern or historical datasets, this correction was not sufficient to allow for reliable age-at-death estimation from ancient human individuals, neither as continuous predictions nor as binary classifications. Predicted ages showed limited consistency with anthropological age estimates, and no CpG window size produced statistically supported age classification. Together, these findings indicate that while correction is a necessary step for mitigating systematic biases inherent to damage-derived methylation inference, it does not fully reconcile the fundamental differences between reconstructed ancient methylomes and the assumptions under which modern epigenetic clock models are trained. Consequently, reliable age-at-death inference from ancient human genomes, even at sequencing depths up to ∼35X, as of now, remains out of reach.

These conclusions must be interpreted in the light of several limitations inherent to biomolecular data. First, the number of ancient individuals available for analysis remains limited, restricting statistical power and the ability to train or robustly evaluate supervised models across age categories. This limitation is particularly relevant for classification and cross-validation analyses, as small and imbalance sample sized increase sensitivity to individual samples and preprocessing choices Second, the precision of damage-derived methylation estimates is constrained by sequencing depth, which limits the reliability of site-specific methylation level estimation and thereby the precision needed for age-at-death estimation at coverage levels typical of ancient genomes. Third, most ancient genomes analyzed in this study were derived from petrous bone, a skeletal element whose inner portion forms early in life and is thought to exhibit limited remodeling and low turnover rates ^38^. As a consequence, methylation patterns recovered from petrous bone may reflect an effectively younger or more homogenized biological age than the true individual age at death. Although the few samples derived from other skeletal elements, such as teeth and long bones, did not yield improved age inference in our dataset, the limited number of such samples precludes strong conclusions.

Taken together, we show that ancient epigenetic age estimation is currently limited by both data availability and methodological constraints. Future progress will hinge on larger and more diverse ancient datasets, systematic comparisons across skeletal tissues and developmental stages, and the development of bone-specific reference methylomes. Closer collaboration with forensic epigenetics and the integration of emerging methylation profiling technologies, such as array-based assays, bisulfite sequencing, or nanopore sequencing may ultimately enable the development of age-prediction models explicitly optimized for ancient DNA and, in the longer term, strengthen demographic reconstructions of past populations and revive palaeodemographic research.

## Acknowledgements

We would like to thank Nicoletta Zedda, Laura Winkelbach, Maxime Brami, Joan Barau, Miguel Andrade, and Daniel Wegmann for the helpful discussions, Katterinne Mendez for assistance, and the GenEvo consortium for support. We thank David Reich for access to unpublished genomes from the Allen Ancient Genome Diversity Project (AAGDP). Parts of this research were conducted using the supercomputer MOGON 2 at Johannes Gutenberg University of Mainz (hpc.uni-mainz.de). This work was funded by the Deutsche Forschungsgemeinschaft (DFG, German Research Foundation) – GRK2526/1 – Projectnr. [407023052]; Y.D. was funded by the ERC Advanced Grant [788616] (‘YMPACT’), awarded to Volker Heyd; L.O. was funded by the ERC Synergy Grant [101071707] (‘Horsepower’).

## STAR METHODS

### METHOD DETAILS

#### Whole genome sequencing data collection

We assembled a dataset of 36 ancient human genomes sequenced to high average depth-of-coverage, which were either publicly available ^39–53^, or available upon request from the Allen Ancient Genome Diversity Project ^54^ (Supplementary Table S1). Three criteria guided sample selection: (i) an average sequencing depth greater than 10X, (ii) reports of USER treatment in library preparation, ensuring reliable DNA methylation inference ^28,29^, and (iii) the availability of an anthropological age estimate, at minimum a categorical assignment. Applying these filters yielded 14 adult and 22 subadult individuals. Since DNA methylation is tissue-specific, we aimed to minimize variability by relying primarily on petrous bone samples. However, due to sample limitations, three genomes were derived from other skeletal elements (Supplementary Table S1) and were still added to our dataset.

#### Ancient DNA processing and methylation inference

Published genomes were obtained from their respective ENA repositories in BAM format, further additional unpublished data for previously published individuals were provided by David Reich’s lab (see Supplementary Table S1 for details). After the removal of PCR duplicates with sambamba ^55^, all genomes were re-aligned around a set of known SNPs and InDels using GATK 3.8 ^56^. Variant detection was performed with ATLAS ^57^ as described in Marchi *et al*. 2022 ^45^.

To infer DNA methylation values, we used the DamMet statistical package (v1.0.4) ^29^, with default parameters in both steps, the “-R” flag to indicate the read groups from the BAM files, excluding any truncated read groups, the “-E” flag to exclude sites with true variants, a maximum CpG window “-N” of 10, 15, 20, and 25, and the “-skip_empty_cpg” parameter.

#### Ancient methylome structure assessment

We evaluated two properties of DNA methylation structure. We first confirmed the expected bimodal distribution of F-values inferred by DamMet by plotting genome-wide methylation profiles per autosome, per sample. We then evaluated whether age-related CpGs – as a representative subset of focal CpG sites – retain coherent methylation patterns across the CpG window sizes used in DamMet. Using four WGBS samples from the ENCODE Project ^58^ (experiments accessions: ENCSR012TGL, ENCSR579AXB, ENCSR307KDA, ENCSR766GKN), we calculated the correlation between the methylation level of each age-related focal CpG and the mean methylation of its neighboring CpGs in windows from 2 to 50 CpGs (step size 2). The correlation trend was used to assess the suitability of CpG windows for DamMet inference.

#### Sex estimation

We inferred molecular sex using DNA methylation patterns along the X chromosome. A panel of ∼4,000 CpGs from the watermelon R package (v.2.8.0) ^37^ was extracted, and principal component analysis (R function prcomp, in R package pROC v.1.18.5) was used to visualize separation between males and females, reflecting sex-specific methylation differences.

#### Implementation of the correction step on ancient data

We implemented the two-step methylation transformation described by Liu et al. (2023). For each CpG site, we constructed predictor variables including the DamMet F-value, its square, sequencing depth, and the interaction between F and depth. A Random Forest (RF) regression model was then trained to predict modern reference methylation values, followed by a linear regression to rescale the RF predictions to match the distribution observed in modern data. The linear regression-adjusted values were used as the final transformed DNA methylation values.

As modern references, we used publicly available methylation datasets generated by the Mammalian Methylation Consortium on the HorvathMammalMethylChip40 array ^17^, specifically GSE184221 and GSE184215 ^18^, with a total of 336 samples. For each CpG site represented in these datasets, we calculated the mean methylation value across all modern samples, which served as the baseline for transformation. Following Liu and colleagues (2023), an exponent parameter (X) was optimized by minimizing prediction error of the RF model, and the optimized value was used to transform the modern reference methylation values. RF model was assessed and tuned using repeated random 90/10% train-test split at the sample level, based on these evaluations, 200 trees were selected for the final model in the ranger package (v0.17.0), with mtry=2. Age-associated CpGs (from the reported four modern epigenetic clocks) were set aside during training, as our goal was to evaluate their performance independent of age association. A custom R script was written to reproduce the transformation workflow of Liu et al. (2023), adapted to our dataset and analysis framework.

#### Application of modern epigenetic clocks

We tested the performance of published epigenetic clocks to reconstruct ancient methylomes, before and after correction. The BBT-APM clock ^23^ and the UniversalPanMammalianClock1 and UniversalPanMammalianClock2 ^18^, where each published model was reimplemented in custom Python scripts following mathematical specifications of the original authors. In addition, the Horvath clock ^12^ was used in R using the methylclock package (v1.8.0) ^16^. Clock performance was assessed by comparing predicted ages to anthropological age estimates using three complimentary metrics. First, association was quantified using a Monte Carlo Spearman correlation approach, in which ages were randomly sampled from the reported anthropological age intervals and Spearman’s rank correlation between predicted and sampled ages was calculated across iterations. Second, we calculated the percentage of predictions falling within the reported anthropological age intervals. Third, for predictions falling outside the interval, we quantified the distance to the nearest interval boundary.

#### Developing an ancient epigenetic clock

An elastic net (EN) regression model was trained to predict chronological age from modern DNA methylation values at a predefined panel of age-associated CpGs corresponding to the union of the CpGs used in the four modern epigenetic clocks evaluated in this study. Modern methylation beta values and sample ages were obtained from the same Mammalian Methylation Consortium datasets described above. DNA methylation matrices were subset to CpGs present in both the age-associated panel and the modern data, standardized by z-score scaling, in which the scaling parameters were later retained for application to ancient data.

Model training was performed using glmnet (v4.1-9) with a Gaussian family. The EN mixing parameter (alpha) was selected by evaluating values from 0 to 1 in 0.1 increments using 10-fold cross-validation (CV), choosing the value minimizing mean absolute error (MAE). A final model was then trained using the selected alpha and 10-fold CV, and coefficients at lambda.min were retained.

For ancient samples, corrected methylation F-values were subset to the panel abovementioned, standardized using the modern scaling parameters, and used to generate point predictions. Prediction uncertainty was quantified by bootstrap resampling across CpGs (B=100). For each bootstrap replicate, CpGs were sampled with replacement, modern methylation values were rescaled on the resampled CpGs, the EN model was refit, and ages were predicted for ancient samples. For each individual, 95% confidence interval (95%-CI) were derived from the empirical distribution of bootstrap predictions.

#### Adult-subadult classification

Binary age classification was performed using EN logistic regression implemented in glmnet with (alpha=0.5) and a probability threshold of 0.5. Individuals were classified as subadults (≤18 years) or adults (>18 years). CpGs were restricted to the predefined age-associated panel abovementioned, and CpGs with zero variance across modern samples were excluded.

Ancient class labels were derived from anthropological age estimates, and individuals with ambiguous age assignment (between 15–25 years) were excluded, and remaining samples were assigned to subadult or adult classes. For each ancient dataset, corrected methylation F-values were reshaped into CpG x sample matrices, missing CpG values were imputed to the corresponding modern CpG mean, and values were scaled using modern centering and scaling parameters.

A baseline classifier was trained on modern samples using 10-fold CV with class-balanced weights and lambda selected as lambda.1se. This model was applied to ancient samples to obtain predicted probabilities and class labels. Performance on labeled ancient samples was evaluated using AUC, accuracy, and balanced accuracy. To assess robustness, a domain-anchored analysis was additionally performed by training classifiers on combined modern and labeled ancient data and evaluating performance using *k*-fold CV on ancient samples only (*k*=5).

